# Reduced interferon antagonism but similar drug sensitivity in Omicron variant compared to Delta variant SARS-CoV-2 isolates

**DOI:** 10.1101/2022.01.03.474773

**Authors:** Denisa Bojkova, Marek Widera, Sandra Ciesek, Mark N. Wass, Martin Michaelis, Jindrich Cinatl

## Abstract

The SARS-CoV-2 Omicron variant is currently causing a large number of infections in many countries. A number of antiviral agents are approved or in clinical testing for the treatment of COVID-19. Despite the high number of mutations in the Omicron variant, we here show that Omicron isolates display similar sensitivity to eight of the most important anti-SARS-CoV-2 drugs and drug candidates (including remdesivir, molnupiravir, and PF-07321332, the active compound in paxlovid), which is of timely relevance for the treatment of the increasing number of Omicron patients. Most importantly, we also found that the Omicron variant displays a reduced capability of antagonising the host cell interferon response. This provides a potential mechanistic explanation for the clinically observed reduced pathogenicity of Omicron variant viruses compared to Delta variant viruses.

---

Omicron (B.1.1.529), is a heavily mutated and highly contagious SARS-CoV-2 variant, which was first detected in southern Africa. It has already replaced the previously dominating Delta variant (B.1.617.2) in some places and is expected to become the dominant SARS-CoV-2 variant in most parts of the world. Protection provided by the current vaccines is substantially reduced against Omicron [1-3]. Moreover, there are many immunocompromised individuals who cannot be effectively protected by vaccines [4]. Hence, antiviral therapies will be essential to protect the most vulnerable individuals from severe COVID-19.

A number of antibody therapies have been approved for use in individuals at a high risk from COVID-19 [5]. Moreover, a range of antiviral small molecule drugs are under investigation or already approved for the treatment of COVID-19. Remdesivir, an intravenous inhibitor of the viral RNA-dependent RNA polymerase (nsp12), was the first antiviral drug to be approved for the treatment of COVID-19 [5,6]. Molnupiravir and PF-07321332 are oral antiviral drugs that are hoped to be able to overcome the issues associated with an intravenous agent [5]. Molnupiravir, a derivative of the broad-spectrum antiviral drug ribavirin, is metabolised into the active compound EIDD-1931, which is incorporated into the complementary RNA strand that is used as a template for the synthesis of viral genomic RNA during replication of the SARS-CoV-2 RNA genome. The incorporation of EIDD-1931 into the template strand causes excessive mutations in the newly produced viral genomes, which affect their functionality in a process called ‘error catastrophe’ or ‘lethal mutagenesis’ [7]. In the UK, molnupiravir is approved and treatment of vulnerable SARS-CoV-2-infected individuals early after diagnosis has been started.

The combination of PF-07321332 (nirmatrelvir) and ritonavir (which reduces PF-07321332 metabolism), also known as paxlovid, has been reported to reduce hospitalisation of SARS-CoV-2-infected individuals in clinical trials [5]. Other antiviral drug candidates for SARS-CoV-2 include the protease inhibitors, camostat, nafamostat, and aprotinin, which inhibit cleavage and activation of the viral spike (S) protein by host cell proteases and, in turn, SARS-CoV-2 entry into host cells [8].

Initial findings suggested that antibody therapies display reduced activity against the Omicron variant [3]. However, it remains unclear whether the mutations associated with the Omicron variant may affect SARS-CoV-2 sensitivity to antiviral drugs. Here, we tested the effects of EIDD-1931, ribavirin, remdesivir, favipravir (an additional RNA-dependent RNA polymerase inhibitor that displayed anti-SARS-CoV-2 activity in phase III clinical trials [9]), PF-07321332, nafamostat, camostat, and aprotinin on the replication of two SARS-CoV-2 Omicron (B.1.1.529) isolates (Omicron 1, Omicron 2, see Suppl. Methods) and one Delta (B.1.167.2) isolate (see Suppl. Methods) [10] in Caco-2 and Calu-3 cells as previously described [11-14].

The Omicron isolates infected fewer cells in Calu-3 and Caco-2 cell cultures when compared with the Delta isolate (Figure 1A, Figure 1B), which is in agreement with previous findings in Calu-3 cells [15] and in the hamster upper respiratory tract [16].

**Figure 1.**
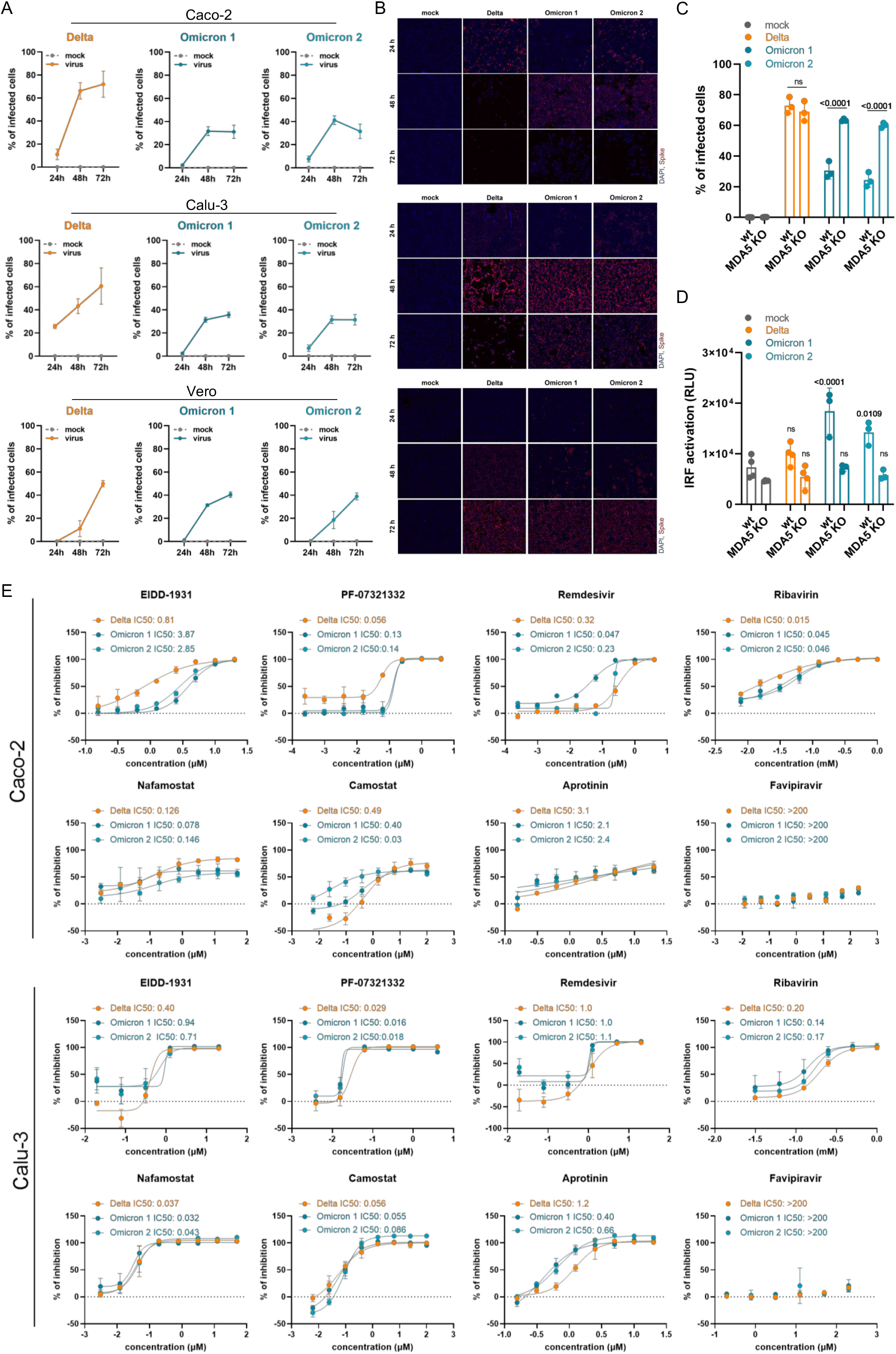
Interferon antagonism and antiviral therapy against novel SARS-CoV-2 variant Omicron. (A) Caco-2 and Calu-3 cells were infected with SARS-CoV-2 variant Delta (GenBank ID: MZ315141), Omicron 1 (GenBank ID: OL800702) and Omicron 2 (GenBank ID: OL800703) at an MOI of 0.01. The number of infected cells at different time points post infection was detected by immunofluorescence staining of the SARS-CoV-2 S protein. Graphs represent mean ± SD of 12 biological replicates. (B) Representative immunofluorescence images of (A) are shown (4x magnification). (C) Virus infection rates in A549-ACE2/TMPRSS2 MDA5-WT (wt) and A549-ACE2/TMPRSS2 MDA5 KO (MDA5 KO) cells 72 h post infection as determined by immunofluorescence staining of the S protein. Graphs represent data of four biological replicates. (D) Induction of IRF transcriptional activity 24 h post infection in a promotor-reporter assay. Graph displays mean ± SD of four biological replicates. (E) Dose dependent effects on SARS-CoV-2 Omicron and Delta variant isolates of selected antiviral compounds. Compounds were added to confluent monolayers and subsequently infected with viral variants at MOI 0.01. The inhibition rate was evaluated 24 h (Caco-2) and 48 h (Calu-3) post infection by staining of the S protein. Graphs depict mean ± SD of three biological replicates. P-values were calculated using two-way ANOVA (C, D). ns – not significant.

However, all three isolates displayed comparable infection patterns in Vero cells (Figure 1A, Figure 1B). In contrast to Caco-2 and Calu-3 cells, Vero cells display a defective interferon response and represent an established model for studying virus replication in an interferon-deficient host cell background [17,18]. Hence, the differences in Omicron virus replication in interferon-competent (Caco-2, Calu-3) and interferon-deficient (Vero) cells suggest that Omicron viruses may be less effective in antagonising cellular interferon signalling than Delta viruses.

In agreement, the Delta isolate displayed superior infection patterns in A549 cells transduced with ACE2 (cellular receptor for the SARS-CoV-2 S protein) and TMPRSS2 (cleaves and activates S) [19], but not in the same cell model with defective interferon signalling due to MDA5 knock-out [20] (Figure 1C). Moreover, the Omicron isolates, but not the Delta isolate, activated interferon signalling as indicated by activation of the interferon response factor (IRF) promotor in A549 cells, which was prevented by MDA5 knock-out (Figure 1D). Taken together, these data show that Omicron viruses are less effective than Delta viruses in antagonising the interferon response in human cells, which may contribute to the lower pathogenicity of the Omicron variant observed in patients [21,22]. Notably, SARS-CoV-2 proteins known to inhibit the host cell interferon response including S, NSP3, NSP6, NSP14, nucleocapsid (N), and membrane (M) are mutated in the Omicron variant [23,24].

Antiviral testing indicated a similar sensitivity of Omicron and Delta isolates to EIDD-1931, PF-07321332, remdesivir, favipravir, ribavirin, nafamostat, camostat, and aprotinin and, hence, to a range of drugs representing different mechanisms of action (Figure 1E). This shows that the mutations in the Omicron variant do not cause substantial changes in the drug sensitivity profiles of the viruses.

With regard to drugs targeting the RNA-dependent RNA polymerase and the replication of the viral genome, this may not come too much as a surprise. Across the replicase-transcriptase complex (nsp7, nsp8, nsp9, nsp10, nsp12, nsp14), only two missense mutations were present in the investigated Omicron isolates, both of which are part of the set of mutations that define the Omicron variant. The RNA-dependent RNA polymerase Nsp12 contains a single change, P323L, which was also present in the Alpha, Beta, and Gamma variants. P323L is far removed from the RNA binding site (Suppl. Figure 1), and would not be expected to impact on RNA implication based on a structural analysis.

One further variant-defining mutation was present in the exonuclease (nsp14), resulting in an I42V change, which is present near the interface site with nsp10. This is a conservative substitution of two small hydrophobic side chains. Structural analysis shows the I42 side chain contacting V40 and N41, which directly contact nsp10 (Suppl. Figure 2). However, this is a minor change that seems unlikely to have a significant impact on the interaction with nsp10 or on antiviral drug activity.

In contrast to our study, which did not detect differences between the sensitivity of Omicron and Delta isolates to TMPRSS2 inhibitors, one previous study had found an Omicron isolate to be less sensitive to camostat than a Delta isolate [15]. Given that this study compared two isolates in one cell line, it is possible that genomic differences between these isolates, which are independent of those defining the Delta and Omicron variant, were responsible for the observed differences. Notably, we detected in Caco-2 cells a 16.3-fold difference between the camostat IC50 for our Delta isolate (0.49µM) compared to the Omicron 2 isolate (0.03µM) (Figure 1E). However, the Omicron 1 isolate displayed a camostat IC50 (0.40µM) very close to that obtained for the Delta isolate, and we did not observe a similar difference in Calu-3 cells (Figure 1E).

Moreover, Omicron mutations are only detected in close vicinity to one of the S cleavage sites. H655Y, N679K, and P681H are close to the 685 furin cleavage site. Among these mutations, only N679K is specific for Omicron (numbering of residues based on the original virus protein sequence). There is no structure for this region of S, because it is a disordered, flexible region. N679K (and P681H) increases the positive charge, but there is no obvious indication that these mutations might affect S cleavage.

In conclusion, our comparison of Omicron and Delta isolates in different cellular models shows that Omicron viruses remain sensitive to a broad range of anti-SARS-CoV-2 drugs and drug candidates with a broad range of mechanisms of action. Moreover, Omicron viruses are less effective in antagonising the host cell interferon response, which may explain why they cause less severe disease [21,22].

## Acknowledgements

This work was supported by the Frankfurter Stiftung für krebskranke Kinder.

## Author contributions

D.B., M.M., and J.C. conceived and designed the study. D.B., M.W., M.N.W., and J.C. performed experiments. All authors analysed data. M.M. wrote the manuscript. D.B., M.N.W., M.M., and J.C. revised the manuscript. All authors have read and approved the final manuscript.

