## Supplemental Methods and Figures for "Reduced interferon antagonism but similar drug sensitivity in Omicron variant compared to Delta variant SARS-CoV-2 isolates"

### Supplementary methods

#### Cell culture

The Caco-2 (DSMZ, Braunschweig, Germany), Vero (DSMZ, Braunschweig, Germany), Calu-3 (ATCC, Manassas, VA, US) were grown at 37 °C in minimal essential medium (MEM) supplemented with 10% fetal bovine serum (FBS), 100 IU/mL of penicillin, and 100 µg/mL of streptomycin. All culture reagents were purchased from Sigma-Aldrich. A Caco-2 subline that was originally established for the cultivation of SARS-CoV and that is also highly permissive to SARS-CoV-2 [11-14] was used for isolation of SARS-CoV-2 variants as well as for antiviral assays. The A549-ACE2/TMPRSS2 (Invivogen) were grown in DMEM supplemented with 2 mM L-glutamine, 4.5 g/l glucose, 10% (v/v) heat-inactivated fetal bovine serum (FBS; 30 min at 56 ºC), PenStrep (100 U/ml-100 µg/ml), 100 µg/ml Normocin, 10 µg/ml of Blasticidin, 10 µg/ml of Blasticidin, 100 µg/ml of Hygromycin, 0.5 µg/ml of Puromycin, and 100 µg/ml of Zeocin. A549-ACE2/TMPRSS2 MDA5 KO cells (Invivogen) were grown in DMEM supplemented with 2 mM L-glutamine, 4.5 g/l glucose, 10% (v/v) heat-inactivated fetal bovine serum (FBS; 30 min at 56 ºC), PenStrep (100 U/ml-100 µg/ml), 100 µg/ml Normocin, 10 µg/ml of Blasticidin, 100 µg/ml of Hygromycin, 0.5 µg/ml of

Puromycin, and 100 µg/ml of Zeocin. All cell lines were regularly authenticated by short tandem repeat (STR) analysis and tested for mycoplasma contamination.

**Virus preparation**

Caco-2 clonal cell subline of human colon carcinoma cell line was used for virus isolation of SARS-CoV-2 variants applied in this study: SARS-CoV-2 Omicron (B.1.1.529) isolates (Omicron 1: FFM-SIM0550/2021, EPI_ISL_6959871, GenBank ID OL800702; Omicron 2: FFM-ZAF0396/2021, EPI_ISL_6959868, GenBank ID OL800703) and one Delta (B.1.167.2) isolate (FFM-IND8424/2021, GenBank ID MZ315141). SARS-CoV-2 stocks used in the experiments had undergone maximum three passages on Caco-2 cells and were stored at –80°C.

**Infection kinetic in cell lines**

The confluent layers of Caco-2, Vero and Calu-3 cells were infected with SARS-CoV-2 variants at MOI 0.01. The number of infected cells was quantified by immunofluorescence staining of spike protein at 24, 48 and 72 h post infection. Briefly, the cells were fixed at indicated times with 3% PFA permeabilized with 0.1 % Triton X-100. Prior to primary antibody labeling, cells were blocked with 5% donkey serum in PBS or 1% BSA and 2% goat serum in PBS for 30 minutes at RT. Spike protein was detected by primary antibody (1:1500, Sinobiological) followed by Alexa Fluor 647 anti-rabbit secondary antibody (1:1000, Invitrogen). The nucleus was labelled using DAPI (1:1000, Thermo Scientific). The images were taken by Spark® Mulitmode microplate reader (TECAN) at 4x magnification.

**IRF induction assay**

A549-ACE2/TMPRSS2 (wt) and A549-ACE2/TMPRSS2 MDA5 KO (MDA5 KO) cells, both containing Lucia luciferase reporter for IRF activity, were seeded in 96-well plates in medium lacking the selection antibiotics. After reaching confluency, the cells were infected with virus at MOI 0.01. The IRF activity was measured in supernatant of infected cells using QUANTI-Luc substrate according manufacturer's protocol 24 h post infection. Briefly, 10 µL of cell culture supernatant were transferred to white cell culture plates and mixed with 50 µL of QUANTI-Luc. The luminescence was measured using Tecan infinite M200 microplate reader (TECAN). The number of infected cells was determined by immunofluorescence staining of spike protein at 72 h post infection.

**Dose-response antiviral assay**

Confluent layers of cells in 96-well plates were treated with decreasing concentration of tested compound and subsequently infected with SARS-CoV-2 at a MOI of 0.01. Evaluation of inhibitory rate was performed by immunohistochemistry of viral spike protein 24 h (Caco-2) or 48 h (Calu-3) post infection.

**Immunocytochemistry of viral antigen**

Cells were fixed with acetone:methanol (40:60) solution and immunostaining was performed using a monoclonal antibody directed against the spike protein of SARS-CoV-2 (1:1500, Sinobiological), which was detected with a peroxidase-conjugated anti-rabbit secondary antibody (1:1,000, Dianova), followed by addition of AEC substrate. The spike positive area was scanned and quantified by the Bioreader® 7000-F-Z-I microplate reader (Biosys). The results are expressed as percentage of inhibition relative to virus control which received no drug.

**Structural analysis**

The defining mutations were identified from https://covariants.org/variants/20I.Alpha.V1 for each variant (link is for alpha). SARS-CoV-2 sequences were obtained from genbank (reference 1af SARS-CoV-2 sequence from Uniprot) and aligned using clustal Omega.

**Statistical analysis**

The results are expressed as the mean ± standard deviation (SD) of number of biological replicates indicated in figure legends. The statistical significance was calculated by two-way ANOVA. GraphPad Prism 9 was used to determine IC50 values.

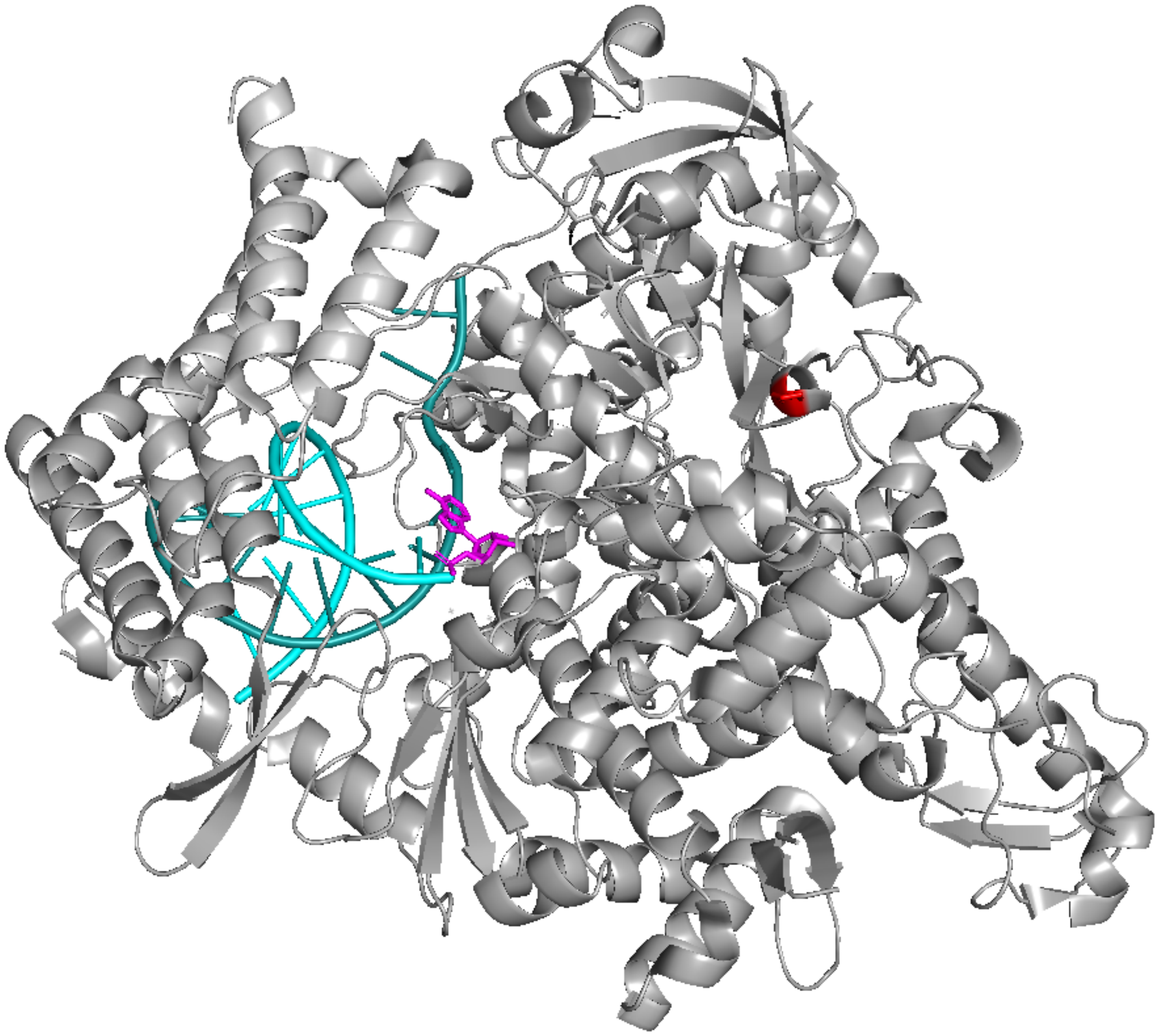

**Supplementary Figure 1** The p.Pro323Leu mutation in the RNA-dependent RNA polymerase. The P323L change is shown in red. RNA is shown in cyan and remdesivir in magenta. Protein structure is protein databank (PDB) code 7bv2. The defining mutations were identified from https://covariants.org/variants/20I.Alpha.V1 for each variant (link is for alpha). The sequences used in this analysis were obtained from the orf1ab sequences from genbank (reference 1af SARS-CoV-2 sequence from Uniprot) and aligned using clustal Omega.

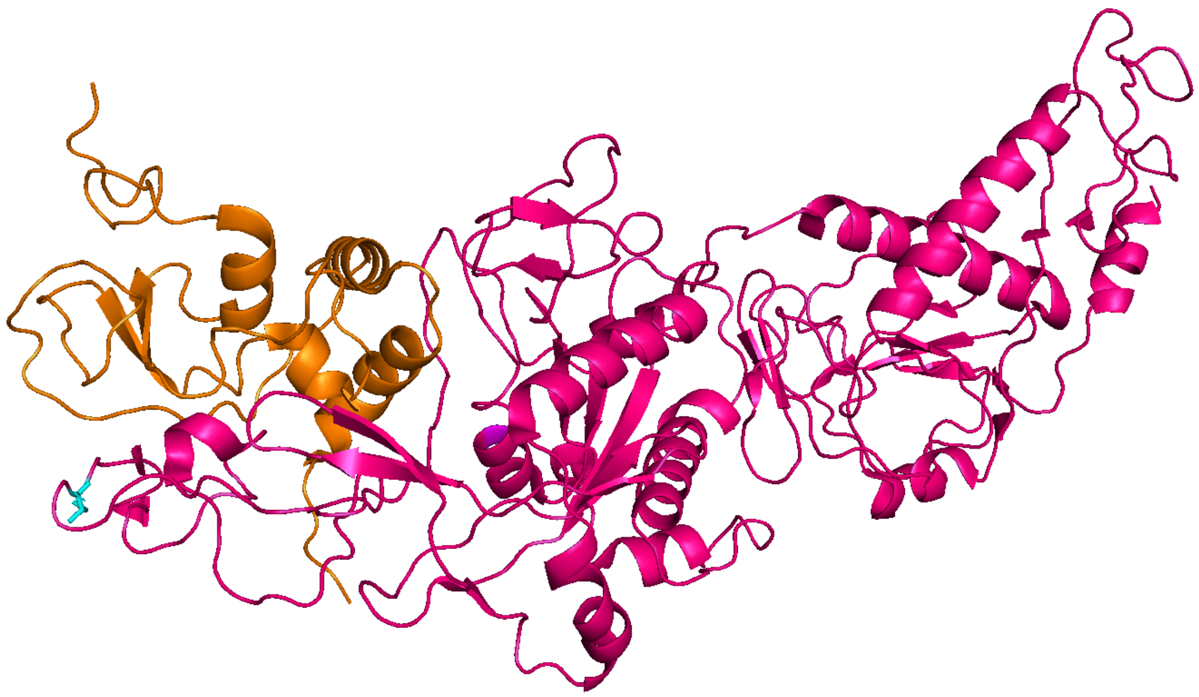

**Supplementary Figure 2**. The I42V change in the exonuclease. The exonuclease (nsp14) is shown in magenta and nsp10 in orange. The I42V change is shown in magenta. Structure shown is from PDB code 7egq. The defining mutations were identified from https://covariants.org/variants/20I.Alpha.V1 for each variant (link is for alpha). The sequences used in this analysis were obtained from the orf1ab sequences from genbank (reference 1af SARS-CoV-2 sequence from Uniprot) and aligned using clustal Omega.
